## Supplementary Material for "Reevaluating the Neural Noise Hypothesis in Dyslexia: Insights from EEG and 7T MRS Biomarkers"

**Table S1.**

*Descriptive Statistics for EEG and MRS Results Separately for the Groups*

|  | DYS | |  | CON | |
| --- | --- | --- | --- | --- | --- |
|  | *M* | *SD* |  | *M* | *SD* |
| *EEG resting state*^a^ |  |  |  |  |  |
| Exponent mean (rest) | 1.54 | 0.14 |  | 1.54 | 0.18 |
| Exponent left IFG (rest) | 1.54 | 0.16 |  | 1.53 | 0.18 |
| Exponent left STS (rest) | 1.50 | 0.18 |  | 1.47 | 0.22 |
| Exponent right IFG (rest) | 1.54 | 0.15 |  | 1.54 | 0.18 |
| Exponent right STS (rest) | 1.48 | 0.18 |  | 1.45 | 0.22 |
| Offset mean (rest) | -10.80 | 0.19 |  | -10.80 | 0.24 |
| Offset left IFG (rest) | -10.72 | 0.34 |  | -10.74 | 0.33 |
| Offset left STS (rest) | -10.97 | 0.38 |  | -10.98 | 0.37 |
| Offset right IFG (rest) | -10.79 | 0.29 |  | -10.81 | 0.32 |
| Offset right STS (rest) | -10.99 | 0.31 |  | -11.04 | 0.36 |
| Beta power left IFG (rest) | 0.48 | 0.18 |  | 0.48 | 0.20 |
| Beta power left STS (rest) | 0.49 | 0.19 |  | 0.48 | 0.21 |
| Beta power right IFG (rest) | 0.49 | 0.18 |  | 0.50 | 0.19 |
| Beta power right STS (rest) | 0.51 | 0.20 |  | 0.50 | 0.21 |
| *EEG language task*^a^ |  |  |  |  |  |
| Exponent mean (task) | 1.55 | 0.15 |  | 1.56 | 0.18 |
| Exponent left IFG (task) | 1.55 | 0.16 |  | 1.55 | 0.19 |
| Exponent left STS (task) | 1.50 | 0.20 |  | 1.47 | 0.21 |
| Exponent right IFG (task) | 1.54 | 0.17 |  | 1.55 | 0.19 |
| Exponent right STS (task) | 1.47 | 0.19 |  | 1.45 | 0.22 |
| Offset mean (task) | -10.67 | 0.25 |  | -10.67 | 0.28 |
| Offset left IFG (task) | -10.58 | 0.39 |  | -10.60 | 0.37 |
| Offset left STS (task) | -10.86 | 0.44 |  | -10.87 | 0.42 |
| Offset right IFG (task) | -10.65 | 0.36 |  | -10.68 | 0.37 |
| Offset right STS (task) | -10.88 | 0.36 |  | -10.94 | 0.41 |
| Beta power left IFG (task)^b^ | 0.50 | 0.23 |  | 0.51 | 0.21 |
| Beta power left STS (task) | 0.54 | 0.24 |  | 0.53 | 0.23 |
| Beta power right IFG (task) | 0.51 | 0.23 |  | 0.52 | 0.21 |
| Beta power right STS (task) | 0.55 | 0.26 |  | 0.55 | 0.23 |
| *MRS* |  |  |  |  |  |
| Glu^c^ | 1.11 | 0.12 |  | 1.07 | 0.10 |
| GABA+^d^ | 0.46 | 0.14 |  | 0.44 | 0.12 |
| Glu/GABA+ ratio^d^ | 2.67 | 0.87 |  | 2.68 | 0.75 |
| Glu/GABA+ imbalance^d^ | 0.27 | 0.11 |  | 0.24 | 0.12 |

*Note.* DYS – dyslexic group; CON – control group; mean – values averaged across all electrodes;

left IFG – values averaged across 3 electrodes corresponding to the left inferior frontal gyrus (F7, FT7, FC5);

left STS – values averaged across 3 electrodes corresponding to the left superior temporal sulcus (T7, TP7, TP9);

right IFG – values averaged across 3 electrodes corresponding to the right inferior frontal gyrus (F8, FT8, FC6);

right STS – values averaged across 3 electrodes corresponding to the right superior temporal sulcus (T8, TP8, TP10);

^a^*n* = 119 (DYS *n* = 59, CON *n* = 60); ^b^*n* = 117 (DYS *n* = 57, CON *n* = 60); ^c^*n* = 50 (DYS *n* = 21, CON *n* = 29);

^d^*n* = 47 (DYS *n* = 20, CON *n* = 27)

**Table S2.**

*Demographic and Behavioral Characteristics of the Subsample of 47 Participants*

|  | DYS  (*n* = 20) | |  | CON  (*n* = 27) | | test | *p* | effect  size | BF_10_ |
| --- | --- | --- | --- | --- | --- | --- | --- | --- | --- |
|  | *M* | *SD* |  | *M* | *SD* |  |  |  |  |
| *Demographics* |  |  |  |  |  |  |  |  |  |
| Sex | 12 F, 8 M | |  | 11 F, 16 M | | χ = 1.71 | .192 | *phi* = -0.19 | 0.81 |
| Age | 19.98 | 3.92 |  | 20.33 | 3.25 | *U* = 261.0 | .846 | *r_rb_* = 0.03 | 0.29 |
| Mother’s education  (years) | 17.48 | 3.68 |  | 16.63 | 2.56 | *U* = 242.5 | .551 | *r_rb_* = -0.10 | 0.36 |
| Father’s education  (years) | 16.74^a^ | 3.46^a^ |  | 16.41 | 3.49 | *U* = 237.0 | .659 | *r_rb_* = -0.08 | 0.31 |
| IQ | 103.80 | 14.30 |  | 113.56 | 9.23 | *t*(30.41) = 2.67 | **.012** | *d* = 0.81 | 6.65 |
| Nonverbal IQ  (scaled score) | 10.45 | 3.14 |  | 12.37 | 2.17 | *U* = 174.5 | **.038** | *r_rb_* = 0.35 | 1.79 |
| ARHQ-PL | 52.90 | 10.78 |  | 24.26 | 6.71 | *U* = 6.0 | **< .001** | *r_rb_* = -0.98 | 1060.02 |
| *Reading and*  *reading-related tasks* |  |  |  |  |  |  |  |  |  |
| Words/min | 110.80 | 19.59 |  | 136.11 | 12.80 | *U* = 74.0 | **< .001*** | *r_rb_* = 0.73 | 108.53 |
| Pseudowords/min | 59.15 | 13.91 |  | 85.59 | 15.83 | *t*(45) = 5.96 | **< .001*** | *d* = 1.76 | >10000 |
| RAN objects (s) | 32.55 | 4.36 |  | 28.56 | 4.46 | *U* = 122.0 | **.001*** | *r_rb_* = -0.55 | 7.92 |
| RAN colors (s) | 36.40 | 4.91 |  | 30.63 | 3.67 | *t*(45) = -4.61 | **< .001*** | *d* = -1.36 | 571.37 |
| RAN digits(s) | 19.35 | 4.00 |  | 16.33 | 2.24 | *U* = 134.0 | **.003*** | *r_rb_* = -0.50 | 3.82 |
| RAN letters (s) | 21.75 | 3.24 |  | 19.52 | 2.53 | *t*(45) = -2.65 | **.011** | *d* = -0.78 | 4.52 |
| Reading comprehension (s) | 63.10 | 18.30 |  | 43.89 | 7.87 | *U* = 82.0 | **< .001*** | *r_rb_* = -0.70 | 65.88 |
| Phoneme deletion  (% correct) | 74.62 | 29.20 |  | 94.16 | 6.61 | *U* = 120.0 | **.001*** | *r_rb_* = 0.56 | 13.46 |
| Spoonerisms phonemes  (% correct) | 52.14 | 38.23 |  | 87.83 | 8.36 | *U* = 106.5 | **< .001*** | *r_rb_* = 0.61 | 17.43 |
| Spoonerisms syllables  (% correct) | 41.67 | 29.86 |  | 77.78 | 20.67 | *U* = 87.5 | **< .001*** | *r_rb_* = 0.68 | 27.77 |
| Orthographic awareness  (accuracy/time) | 0.35 | 0.16 |  | 0.54 | 0.13 | *t*(45) = 4.26 | **< .001*** | *d* = 1.26 | 215.19 |
| Perception speed (sten score) | 2.95 | 1.79 |  | 4.33 | 1.64 | *U* = 126.0 | **.002*** | *r_rb_* = 0.53 | 4.49 |
| Digits forward | 5.65 | 1.79 |  | 7.04 | 2.03 | *U* = 158.0 | **.014** | *r_rb_* = 0.42 | 2.13 |
| Digits backward | 4.95 | 1.73 |  | 7.48 | 2.06 | *t*(45) = 4.45 | **< .001*** | *d* = 1.31 | 355.64 |

*Note.* DYS – dyslexic group; CON – control group; F – females, M – males. BF_10_ – Bayes Factor indicating ratio of the likelihood of an alternative hypothesis (H1) to a null hypothesis (H0). ARHQ-PL – Polish version of the Adult Reading History Questionnaire. RAN – rapid automatized naming. Non-parametric Mann-Whitney test was performed when assumption of normal distribution was violated. *r_rb_* – rank biserial correlation provided as an effect size parameter for Mann-Whitney test. Boldface indicates statistical significance at *p* < .05 level (uncorrected).

*Significance after Bonferroni correction for 14 planned comparisons for reading and reading-related tasks;

^a^*n* = 19 (one participant did not provide information about the father’s education)

**EEG Data – Frontal and Temporal Electrodes**

**Beta (14-30 Hz) aperiodic-adjusted**

*Beta center frequency*

There was a significant effect of condition (*F*(1,115) = 6.12, *p* = .015, η^2^_p_ = .051, BF_incl_ = 2.94) with post-hoc comparison indicating that the beta peak was at higher frequencies at rest (*M* = 19.86, *SD* = 2.64) than during the language task (*M* = 19.44, *SD* = 2.48, *p_corr_* = .015). There was also a significant interaction between condition and region, although the Bayes Factor did not provide evidence for either inclusion or exclusion (*F*(1,115) = 5.96, *p* = .016, η^2^_p_ = .049, BF_incl_ = 1.52). Post-hoc comparisons indicated that the beta peak was at higher frequencies at rest than during the language task in the frontal region (*M*_rest_ = 20.03, *SD*_rest_ = 2.79, *M*_task_ = 19.43, *SD*_task_ = 2.51, *p_corr_* = .002), while this difference was not significant in the temporal region (*M*_rest_ = 19.68, *SD*_rest_ = 2.76, *M*_task_ = 19.45, *SD*_task_ = 2.70, *p_corr_* = .207). Furthermore, at rest, the beta peak was at higher frequencies in the frontal as compared to the temporal region (*p_corr_* = .028), while this difference was not significant during the language task (*p_corr_* = .878). The effect of group (*F*(1,115) = 0.02, *p* = .896, η^2^_p_ = .000, BF_incl_ = 0.001) was not significant and Bayes Factor indicated against including it in the model. Any other effects of interactions were not significant and Bayesian statistics indicated against including these factors in the model or did not provide evidence for either inclusion or exclusion.

*Beta bandwidth*

The interaction between region, hemisphere, and group was not significant, although Bayesian statistics indicated in favor of including it in the model (*F*(1,115) = 1.92, *p* = .169, η^2^_p_ = .016, BF_incl_ = 389.67). The effect of group (*F*(1,115) = 0.39, *p* = .532, η^2^_p_ = .003, BF_incl_ = 0.60) was not significant while Bayes Factor did not provide evidence for either inclusion or exclusion. Any other effects of interactions were not significant and Bayesian statistics indicated against including them in the model or did not provide evidence for either inclusion or exclusion. Since Bayes Factor suggested the inclusion of the region*hemisphere*group interaction in the model, we further conducted Bayesian *t*-tests to determine whether this was driven by differences between control and dyslexic groups. The results, however, supported the null hypothesis in both the left (*M*_DYS_ = 7.19, *SD*_DYS_ = 2.64, *M*_CON_ = 6.96, *SD*_CON_ = 2.84, BF_10_ = 0.22) and right hemisphere in the frontal region (*M*_DYS_ = 6.93, *SD*_DYS_ = 2.86, *M*_CON_ = 7.07, *SD*_CON_ = 2.80, BF_10_ = 0.20) as well as in the left hemisphere in the temporal region (*M*_DYS_ = 7.32, *SD*_DYS_ = 2.57, *M*_CON_ = 6.86, *SD*_CON_ = 2.72, BF_10_ = 0.29). The results in the right hemisphere in the temporal region were inconclusive (*M*_DYS_ = 7.09, *SD*_DYS_ = 2.25, *M*_CON_ = 6.36, *SD*_CON_ = 2.61, BF_10_ = 0.66).

**Alpha (7-14 Hz) aperiodic-adjusted**

For these analyses, the sample size was 112 (DYS *n* = 56, CON *n* = 56), since in 7 participants the algorithm did not find the alpha peak above the aperiodic component in selected electrodes.

*Alpha power*

There was a significant effect of condition (*F*(1,110) = 63.47, *p* < .001, η^2^_p_ = .366, BF_incl_ > 10000) with post-hoc comparison indicating that the alpha power was greater during the language task (*M* = 1.21, *SD* = 0.47) than at rest (*M* = 0.99, *SD* = 0.39, *p_corr_* < .001). There were also significant effects of hemisphere (*F*(1,110) = 13.84, *p* < .001, η^2^_p_ = .112, BF_incl_ = 76.81) and region (*F*(1,110) = 6.34, *p* = .013, η^2^_p_ = .054, BF_incl_ = 2.98). For the main effect of hemisphere, post-hoc comparison indicated that alpha power was greater in the right (*M* = 1.11, *SD* = 0.41) as compared to the left hemisphere (*M* = 1.09, *SD* = 0.42, *p_corr_* < .001), while for the main effect of region, post-hoc comparison indicated that the alpha power was greater in the temporal (*M* = 1.11, *SD* = 0.41) as compared to the frontal region (*M* = 1.09, *SD* = 0.42, *p_corr_* = .013). Furthermore, there were significant interactions between condition, region, and group (*F*(1,110) = 4.78, *p* = .031, η^2^_p_ = .042, BF_incl_ = 64.84) as well as between hemisphere and region (*F*(1,110) = 4.35, *p* = .039, η^2^_p_ = .038, BF_incl_ = 0.92) although Bayes Factor did not provide evidence for either inclusion or exclusion the hemisphere*region interaction. For the condition*region*group interaction, post-hoc comparisons indicated that in both groups and in both regions, alpha power was greater in the language task than at rest (for all comparisons *p_corr_* < .001). Furthermore, in the control group during resting state condition, alpha power was greater in the temporal (*M* = 0.99, *SD* = 0.36) as compared to the frontal region (*M* = 0.95, *SD* = 0.38, *p_corr_* = .003), while any other comparisons were not significant. For the hemisphere*region interaction, post-hoc comparisons indicated that greater alpha power in the temporal as compared to the frontal region was significant in the right (*M*_frontal_ = 1.10, *SD*_frontal_ = 0.42, *M*_temporal_ = 1.13, *SD*_temporal_ = 0.40, *p_corr_* = .001), while not in the left hemisphere (*M*_frontal_ = 1.08, *SD*_frontal_ = 0.42, *M*_temporal_ = 1.09, *SD*_temporal_ = 0.42, *p_corr_* = .386). Also, in the temporal region, greater alpha power was found in the right than in the left hemisphere (*p_corr_* < .001), while the difference between hemispheres was not significant in the frontal region (*p_corr_* = .110). The effect of group (*F*(1,110) = 0.27, *p* = .607, η^2^_p_ = .002, BF_incl_ = 0.02) was not significant and Bayes Factor indicated against including it in the model. Any other interactions were not significant and Bayesian statistics indicated against including them in the model or did not provide evidence for either inclusion or exclusion.

*Alpha center frequency*

There was a significant effect of condition (*F*(1,110) = 15.24, *p* < .001, η^2^_p_ = .122, BF_incl_ = 144.27) with post-hoc comparison indicating that alpha peak was at lower frequencies at rest (*M* = 10.51, *SD* = 0.98) than during the language task (*M* = 10.73, *SD* = 0.94, *p_corr_* < .001). There were also significant interactions between condition and hemisphere (*F*(1,110) = 9.99, *p* = .002, η^2^_p_ = .083, BF_incl_ = 14.42), as well as between condition and region (*F*(1,110) = 4.28, *p* = .041, η^2^_p_ = .037, BF_incl_ = 0.82), although Bayes Factor did not provide evidence for either including or excluding condition*region interaction. For the condition*hemisphere interaction, post-hoc comparisons indicated that the alpha peak was at lower frequencies at rest than during the language task both in the left (*M*_rest_ = 10.59, *SD*_rest_ = 1.04, *M*_task_ = 10.72, *SD*_task_ = 0.95, *p_corr_* = .048) and in the right hemisphere (*M*_rest_ = 10.42, *SD*_rest_ = 1.03, *M*_task_ = 10.75, *SD*_task_ = 0.95, *p_corr_* < .001). Furthermore, in the resting state condition, the alpha peak was at lower frequencies in the right as compared to the left hemisphere (*p_corr_* = .008), while the difference between hemispheres was not significant during the language task (*p_corr_* = .334). For the condition*region interaction, post-hoc comparisons indicated that alpha peak was at lower frequencies at rest than during the language task both in the temporal (*M*_rest_ = 10.48, *SD*_rest_ = 0.98, *M*_task_ = 10.76, *SD*_task_ = 0.96, *p_corr_* < .001) and in the frontal region (*M*_rest_ = 10.53, *SD*_rest_ = 1.02, *M*_task_ = 10.71, *SD*_task_ = 0.97, *p_corr_* = .008), while the difference between frontal and temporal regions was not significant either at rest (*p_corr_* = .128) or during the language task (*p_corr_* = .288). The effect of group (*F*(1,110) = 1.55, *p* = .216, η^2^_p_ = .014, BF_incl_ = 0.70) was not significant while Bayes Factor did not provide evidence for either inclusion or exclusion. Any other interactions were not significant and Bayesian statistics indicated against including them in the model or did not provide evidence for either inclusion or exclusion.

*Alpha bandwidth*

The analysis revealed a significant effect of condition (*F*(1,110) = 6.21, *p* = .014, η^2^_p_ = .053, BF_incl_ = 3.06) with post-hoc comparison indicating that the alpha peak was wider at rest (*M* = 3.18, *SD* = 1.25) than during the language task (*M* = 2.91, *SD* = 0.94, *p_corr_* = .014). There was also a significant effect of region, although Bayesian statistics did not provide evidence for either inclusion or exclusion (*F*(1,110) = 5.42, *p* = .022, η^2^_p_ = .047, BF_incl_ = 1.64). Post-hoc comparison indicated that the alpha peak was wider in the temporal (*M* = 3.12, *SD* = 0.94) as compared to the frontal region (*M* = 2.97, *SD* = 1.04, *p_corr_* = .022). There were also significant interactions between region and condition (*F*(1,110) = 7.33, *p* = .008, η^2^_p_ = .062, BF_incl_ = 4.15) as well as between region and group (*F*(1,110) = 5.59, *p* = .020, η^2^_p_ = .048, BF_incl_ = 0.38), although Bayes Factor did not provide evidence for either including or excluding region*group interaction. For the region*condition interaction, post-hoc comparisons indicated that the alpha peak was wider in the resting state condition as compared to the language task in the frontal region (*M*_rest_ = 3.18, *SD*_rest_ = 1.45, *M*_task_ = 2.77, *SD*_task_ = 0.99, *p_corr_* = .001), while this difference was not significant in the temporal region (*M*_rest_ = 3.19, *SD*_rest_ = 1.22, *M*_task_ = 3.04, *SD*_task_ = 1.03, *p_corr_* = .225). Furthermore, during the language task, the alpha peak was wider in the temporal than in the frontal region (*p_corr_* < .001), while the difference between regions was not significant during the resting state condition (*p_corr_* = .932). For the region*group interaction, post-hoc comparisons indicated that in the dyslexic group, the alpha peak was wider in the temporal as compared to the frontal region (*M*_frontal_ = 2.85, *SD*_frontal_ = 1.09, *M*_temporal_ = 3.14, *SD*_temporal_ = 1.01, *p_corr_* = .001), while this difference was not significant in the control group (*M*_frontal_ = 3.10, *SD*_frontal_ = 0.98, *M*_temporal_ = 3.09, *SD*_temporal_ = 0.87, *p_corr_* = .980). The difference between dyslexic and control groups was not significant either in the frontal (*p_corr_* = .214) or in the temporal region (*p_corr_* = .810). The interaction between region, hemisphere, and condition was not significant, although Bayesian statistics indicated in favor of including it in the model (*F*(1,110) = 1.54, *p* = .217, η^2^_p_ = .014, BF_incl_ = 5.96). The effect of group (*F*(1,110) = 0.33, *p* = .569, η^2^_p_ = .003, BF_incl_ = 0.05) was not significant and Bayes Factor indicated against including it in the model. Any other interactions were not significant and Bayesian statistics indicated against including them in the model or did not provide evidence for either inclusion or exclusion**.**

**EEG Data – Parieto-Occipital Electrodes**

Following the previous study, which revealed differences in aperiodic and periodic components between dyslexic and control groups in the parieto-occipital region (Turri et al., 2023), we conducted additional analyses using the same cluster of electrodes from the left (PO7, PO3, O1) and the right hemisphere (PO8, PO4, O2). For the exponent and offset, we employed a 2x2x2 (group, condition, hemisphere) repeated measures ANOVA with age included as a covariate. For the beta and alpha bands results, we used a similar model but without the effect of age included as a covariate.

**Exponent**

The analysis revealed significant effects of age (*F*(1,116) = 5.22, *p* = .024, η^2^_p_ = .043, BF_incl_ = 2.07) and hemisphere (*F*(1,116) = 6.37, *p* = .013, η^2^_p_ = .052, BF_incl_ > 10000) with post-hoc comparison indicating that the exponent was lower in the left (*M* = 1.46, *SD* = 0.21) as compared to the right hemisphere (*M* = 1.53, *SD* = 0.19, *p_corr_* < .001). The effect of group was not significant *F*(1,116) = 0.07, *p* = .786, η^2^_p_ = .001, BF_incl_ = 0.65) although Bayes Factor did not provide evidence for either inclusion or exclusion. Any other effects or interactions were not significant and Bayesian statistics indicated against including them in the model or did not provide evidence for either inclusion or exclusion.

**Offset**

There were significant effects of hemisphere (*F*(1,116) = 15.20, *p* < .001, η^2^_p_ = .116, BF_incl_ > 10000) and condition (*F*(1,116) = 8.70, *p* = .004, η^2^_p_ = .070, BF_incl_ > 10000) with post-hoc comparisons indicating that the offset was lower in the left (*M* = -11.19, *SD* = 0.52) as compared to the right hemisphere (*M* = -10.73, *SD* = 0.27, *p_corr_* < .001), and at rest (*M* = -11.03, *SD* = 0.35) than during the language task (*M* = -10.90, *SD* = 0.39, *p_corr_* < .001). The interaction between condition and hemisphere (*F*(1,116) = 0.13, *p* = .725, η^2^_p_ = .001, BF_incl_ = 31.62) was not significant although Bayes Factor indicated in favor of including it in the model. The effect of group (*F*(1,116) = 0.08, *p* = .781, η^2^_p_ = .001, BF_incl_ = 0.04) was not significant and Bayes Factor indicated against including it in the model. Any other effects or interactions were not significant and Bayesian statistics indicated against including them in the model or did not provide evidence for either inclusion or exclusion.

**Beta (14-30 Hz) aperiodic-adjusted**

*Beta power*

The analysis revealed significant effects of hemisphere (*F*(1,117) = 18.74, *p* < .001, η^2^_p_ = .138, BF_incl_ = 612.30) and condition (*F*(1,117) = 24.05, *p* < .001, η^2^_p_ = .170, BF_incl_ = 4545.40). For the main effect of hemisphere, post-hoc comparison indicated that the beta power was greater in the right (*M* = 0.56, *SD* = 0.19) as compared to the left hemisphere (*M* = 0.53, *SD* = 0.18, *p_corr_* < .001), while for the main effect of condition, post-hoc comparison indicated that the beta power was greater during the language task (*M* = 0.57, *SD* = 0.21) than at rest (*M* = 0.51, *SD* = 0.18, *p_corr_* < .001). The effect of group was not significant (*F*(1,117) = 0.06, *p* = .841, η^2^_p_ = .000, BF_incl_ = 0.55), although Bayes Factor did not provide evidence for either inclusion or exclusion. Any other interactions were not significant and Bayesian statistics indicated against including them in the model or did not provide evidence for either inclusion or exclusion.

*Beta center frequency*

The analysis revealed significant interactions between group and hemisphere *F*(1,117) = 5.10, *p* = .026, η^2^_p_ = .042, BF_incl_ = 1.74), as well as between group, hemisphere, and condition *F*(1,117) = 4.15, *p* = .044, η^2^_p_ = .034, BF_incl_ = 1.89), although Bayes Factor did not provide evidence for either inclusion or exclusion. For the group*hemisphere interaction, post-hoc comparisons did not reveal any significant differences. For the group*hemisphere*condition interaction, post-hoc comparisons indicated that in the dyslexic group during the resting state condition, beta peak was at lower frequencies in the right (*M* = 18.51, *SD* = 1.95) as compared to the left hemisphere (*M* = 19.07, *SD* = 2.24, *p_corr_* = .026), while any other comparisons were not significant. The effect of group was not significant (*F*(1,117) = 0.20, *p* = .659, η^2^_p_ = .002, BF_incl_ = 0.37), although Bayes Factor did not provide evidence for either inclusion or exclusion. Any other effects or interactions were not significant and Bayesian statistics indicated against including them in the model.

*Beta bandwidth*

The effect of group was not significant (*F*(1,117) = 0.02, *p* = .890, η^2^_p_ = .000, BF_incl_ = 0.19) and Bayes Factor indicated against including it in the model. Any other effects or interactions were not significant and Bayesian statistics indicated against including them in the model or did not provide evidence for either inclusion or exclusion.

**Alpha (7-14 Hz) aperiodic-adjusted**

For these analyses, the sample size was 117 (DYS *n* = 59, CON *n* = 58), since in 2 participants the algorithm did not find the alpha peak above the aperiodic component in selected electrodes.

*Alpha power*

There were significant effects of hemisphere (*F*(1,115) = 63.01, *p* < .001, η^2^_p_ = .354, BF_incl_ > 10000) and condition (*F*(1,115) = 93.58, *p* < .001, η^2^_p_ = .449, BF_incl_ > 10000). For the main effect of hemisphere, post-hoc comparison indicated that the alpha power was greater in the right (*M* = 1.30, *SD* = 0.36) as compared to the left hemisphere (*M* = 1.22, *SD* = 0.34, *p_corr_* < .001), while for the main effect of condition, post-hoc comparison indicated that the alpha power was greater during the language task (*M* = 1.38, *SD* = 0.36) than at rest (*M* = 1.15, *SD* = 0.38, *p_corr_* < .001). There were also significant interactions between group and hemisphere (*F*(1,115) = 5.25, *p* = .024, η^2^_p_ = .044, BF_incl_ = 2.26), as well as between hemisphere and condition (*F*(1,115) = 4.01, *p* = .048, η^2^_p_ = .034, BF_incl_ = 1.36), however Bayes Factor did not provide the evidence for either inclusion or exclusion. For the group*hemisphere interactions, post-hoc comparisons indicated that greater alpha power was in the right as compared to the left hemisphere both in the dyslexic (*M*_left_ = 1.20, *SD*_left_ = 0.35, *M*_right_ = 1.31, *SD*_right_ = 0.36, *p_corr_* < .001) and in the control group (*M*_left_ = 1.24, *SD*_left_ = 0.33, *M*_right_ = 1.30, *SD*_right_ = 0.36, *p_corr_* < .001), while the difference between the dyslexic and control group was not significant either in the left (*p_corr_* < .497), or in the right hemisphere (*p_corr_* < .926). For the hemisphere*condition interaction, post-hoc comparisons indicated that the alpha power was greater in the right as compared to the left hemisphere both at rest and during the language task (all comparisons *p_corr_* < .001), and that the alpha power was greater during the task than at rest both in the left and in the right hemisphere (all comparisons *p_corr_* < .001). The effect of group was not significant (*F*(1,115) = 0.08, *p* = .776, η^2^_p_ = .001, BF_incl_ = 0.56), although Bayes Factor did not provide evidence for either inclusion or exclusion. Any other interactions were not significant and Bayesian statistics indicated against including them in the model or did not provide evidence for either inclusion or exclusion.

*Alpha center frequency*

There was a significant effect of condition (*F*(1,115) = 92.36, *p* < .001, η^2^_p_ = .445, BF_incl_ > 10000) with post-hoc comparison indicating that the alpha peak was at lower frequencies at rest (*M* = 10.44, *SD* = 0.95) than during the language task (*M* = 10.87, *SD* = 0.93, *p_corr_* < .001). The effect of group was not significant (*F*(1,115) = 2.94, *p* = .089, η^2^_p_ = .025, BF_incl_ = 0.60), although Bayes Factor did not provide evidence for either inclusion or exclusion. Any other effects or interactions were not significant and Bayesian statistics indicated against including them in the model or did not provide evidence for either inclusion or exclusion.

*Alpha bandwidth*

The effect of group was not significant (*F*(1,115) = 0.01, *p* = .923, η^2^_p_ = .000, BF_incl_ = 0.36), although Bayes Factor did not provide evidence for either inclusion or exclusion. Any other effects or interactions were not significant and Bayesian statistics indicated against including them in the model or did not provide evidence for either inclusion or exclusion.

**Table S3.**

*Zero-order Correlations Between MRS and EEG Biomarkers of Excitatory-Inhibitory Balance*

| Variable | 1.  *r*  (BF_10_) | 2. | 3. | 4. | 5. | 6. | 7. | 8. |
| --- | --- | --- | --- | --- | --- | --- | --- | --- |
| *EEG resting state* |  |  |  |  |  |  |  |  |
| 1. Glu | – |  |  |  |  |  |  |  |
| 2. GABA+ | .40**_a_  (8.08) | – |  |  |  |  |  |  |
| 3. Glu/GABA+ ratio | -.11_a_  (0.24) | -.90***_a_  (>10000) | – |  |  |  |  |  |
| 4. Glu/GABA+ imbalance | .16_a_  (0.32) | .33*_a_  (2.14) | -.19_a_  (0.42) | – |  |  |  |  |
| 5. Exponent  mean (rest) | .13 _b_  (0.25) | .10_a_  (0.23) | -.11_a_  (0.24) | .31*_a_  (1.58) | – |  |  |  |
| 6. Offset  mean (rest) | .10_b_  (0.23) | .17_a_  (0.35) | -.15_a_  (0.30) | .26_a_  (0.79) | .70***_c_  (>10000) | – |  |  |
| 7. Exponent  left STS (rest) | .04_b_  (0.18) | .09_a_  (0.22) | -.08_a_  (0.21) | .37*_a_  (4.06) | .70***_c_  (>10000) | .52***_c_  (>10000) | – |  |
| 8. Offset  left STS (rest) | -.09_b_  (0.21) | .01_a_  (0.18) | .00_a_  (0.18) | .21_a_  (0.48) | .24**_c_  (3.92) | .54***_c_  (>10000) | .70***_c_  (>10000) | – |
| 9. Beta power  left STS (rest) | -.06_b_  (0.19) | .22_a_  (0.53) | -.28_a_  (1.04) | .03_a_  (0.19) | .19*_c_  (0.85) | .20*_c_  (1.29) | .50***_c_  (>10000) | .56***_c_  (>10000) |
| *EEG language task* |  |  |  |  |  |  |  |  |
| 5. Exponent  mean (task) | .14_b_  (0.27) | .15_a_  (0.29) | -.14_a_  (0.28) | .24_a_  (0.64) | – |  |  |  |
| 6. Offset  mean (task) | .10_b_  (0.23) | .19_a_  (0.41) | -.16_a_  (0.31) | .23_a_  (0.59) | .75***_c_  (>10000) | – |  |  |
| 7. Exponent  left STS (task) | .05_b_  (0.19) | .09_a_  (0.22) | -.09_a_  (0.21) | .24_a_  (0.64) | .69***_c_  (>10000) | .58***_c_  (>10000) | – |  |
| 8. Offset  left STS (task) | -.07_b_  (0.20) | .03_a_  (0.19) | -.01_a_  (0.18) | .18_a_  (0.38) | .36***_c_  (375.45) | .62***_c_  (>10000) | .80***_c_  (>10000) | – |
| 9. Beta power  left STS (task) | -.07_b_  (0.20) | .22_a_  (0.52) | -.25_a_  (0.77) | -.01_a_  (0.18) | .07_c_  (0.15) | .15_c_  (0.43) | .47***_c_  (>10000) | .59***_c_  (>10000) |

*Note. r* – Pearson’s correlation coefficient; BF_10_ – Bayes Factor indicating ratio of the likelihood of an alternative hypothesis (H1) to a null hypothesis (H0); mean – values averaged across all electrodes; left STS – values averaged across 3 electrodes corresponding to the left superior temporal sulcus (T7, TP7, TP9).

****p* < .001 (uncorrected); ***p* < .01 (uncorrected); **p* < .05 (uncorrected);

^a^*n* = 47; ^b^*n* = 50; ^c^*n* = 119

**Table S4.**

*Zero-order Correlations Between Reading, Phonological Awareness, Rapid Automatized Naming, Multisensory Integration and Biomarkers of Excitatory-Inhibitory Balance*

| Variable | 1.  *r*  (BF_10_) | 2. | 3. | 4. |
| --- | --- | --- | --- | --- |
| *EEG resting state* |  |  |  |  |
| 1. Reading | – |  |  |  |
| 2. Phonological awareness | .62***_c_  (>10000) | – |  |  |
| 3. RAN | .73***_c_  (>10000) | .50***_c_  (>10000) | – |  |
| 4. Multisensory integration | .24*_d_  (1.44) | .33**_d_  (16.94) | .08_d_  (0.17) | – |
| 5. Glu | -.10_b_  (0.22) | -.16_b_  (0.32) | -.08_b_  (0.20) | -.08_b_  (0.21) |
| 6. GABA+ | -.17_a_  (0.35) | -.02_a_  (0.18) | -.19_a_  (0.41) | .24_a_  (0.62) |
| 7. Glu/GABA+ ratio | .03_a_  (0.19) | -.02_a_  (0.18) | .08_a_  (0.21) | -.31*_a_  (1.62) |
| 8. Glu/GABA+ imbalance | -.21_a_  (0.47) | -.06_a_  (0.20) | -.19_a_  (0.40) | .13_a_  (0.27) |
| 9. Exponent mean  (rest) | -.13_c_  (0.30) | .05_c_  (0.13) | -.08_c_  (0.17) | -.05_d_  (0.15) |
| 10. Offset mean  (rest) | -.03_c_  (0.12) | .04_c_  (0.13) | -.02_c_  (0.12) | -.01_d_  (0.14) |
| 11. Exponent left STS (rest) | -.14_c_  (0.37) | -.01_c_  (0.12) | -.07_c_  (0.16) | -.15_d_  (0.33) |
| 12. Offset left STS  (rest) | .03_c_  (0.12) | .09_c_  (0.18) | .03_c_  (0.12) | -.07_d_  (0.17) |
| 13. Beta power left STS (rest) | .03_c_  (0.12) | .22*_c_  (1.96) | -.04_c_  (0.12) | .04_d_  (0.14) |
| *EEG language task* |  |  |  |  |
| 9. Exponent mean  (task) | -.13_c_  (0.32) | .06_c_  (0.14) | -.14_c_  (0.34) | -.09_d_  (0.19) |
| 10. Offset mean  (task) | -.05_c_  (0.13) | .04_c_  (0.12) | -.05_c_  (0.13) | -.01_d_  (0.13) |
| 11. Exponent left STS (task) | -.11_c_  (0.23) | .01_c_  (0.12) | -.11_c_  (0.24) | -.17_d_  (0.48) |
| 12. Offset left STS  (task) | .04_c_  (0.12) | .09_c_  (0.18) | .01_c_  (0.12) | -.07_d_  (0.16) |
| 13. Beta power left STS (task) | .05_c_  (0.13) | .21*_c_  (1.61) | .02_c_  (0.12) | .11_d_  (0.22) |

*Note. r* – Pearson’s correlation coefficient; BF_10_ – Bayes Factor indicating ratio of the likelihood of an alternative hypothesis (H1) to a null hypothesis (H0); mean – values averaged across all electrodes; left STS – values averaged across 3 electrodes corresponding to the left superior temporal sulcus (T7, TP7, TP9).

****p* < .001 (uncorrected); ***p* < .01 (uncorrected); **p* < .05 (uncorrected);

^a^*n* = 47; ^b^*n* = 50; ^c^*n* = 119; ^d^*n* = 87

**fMRI Data**

Functional data acquired from two participants with a 12-channel radiofrequency coil were acquired using whole-brain echo planar imaging sequence (TE = 28ms, TR = 2500 ms, flip angle FA = 80°, FOV = 216 mm, matrix size = 72x72, 42 axial slices 3 mm thick, 3x3 mm in-plane resolution).

*Group-level analysis*

For this analysis, the following preprocessing steps were conducted: 1) realignment of all functional images to the participant’s mean, 2) coregistration of T1-weighted images to functional images for each subject, 3) segmentation of coregistered anatomical images, 4) normalization of functional images, and 5) smoothing of functional data with a 6 mm isotropic Gaussian kernel*.* We also used ART (artifact detection tools with default options) to add movement regressors and reject volumes identified as motion outliers.

In the second-level analysis, we conducted one-sample *t*-tests to examine activation maps separately for visual (words > false fonts) and auditory runs (words > backward) within each group (dyslexic, control). We also employed paired *t*-tests to assess activations for both visual and auditory runs (logical AND conjunction), examining them separately for the groups. Finally, a flexible factorial model was utilized with the main factor of subject and an interaction factor between group and condition to investigate the effect of group in both visual and auditory runs. We reported the results at *p* < .001 height threshold corrected for multiple comparisons using *p* < .05 FWE cluster threshold. The anatomical regions were labeled based on the AAL3 atlas (Rolls et al., 2020). The results are reported for 50 participants – 21 in the dyslexic (12 females, 9 males) and 29 in the control group (13 females, 16 males) reflecting the sample size for Glu results from the MRS.

**Table S5.** *One Sample T-Tests Separately for Visual (Words > False Fonts) and Auditory Runs (Words > Words Backward) Within Control (CON) and Dyslexic Groups (DYS)*

| Brain regions | Hemisphere | Peak of cluster coordinates | | | *t-*value | Number of voxels |
| --- | --- | --- | --- | --- | --- | --- |
|  |  | x | y | z |  |  |
| **CON (*n* = 29) visual runs** (FWEc = 127) | | | | | | |
| middle temporal gyrus, superior temporal gyrus, inferior frontal gyrus (pars triangularis, orbitalis, opercularis), supramarginal gyrus, temporal pole (superior & middle temporal gyri), postcentral gyrus, hippocampus, posterior orbital gyrus, amygdala, parahippocampal gyrus, inferior parietal gyrus, rolandic operculum, lateral orbital gyrus, pallidum, anterior orbital gyrus, inferior frontal gyrus, putamen, middle frontal gyrus, insula, angular gyrus | L | -62 | -54 | 10 | 9.78 | 5360 |
| precuneus, posterior cingulate gyrus, middle cingulate & paracingulate gyri, calcarine fissure, cuneus | L/R | -4 | -52 | 20 | 7.17 | 957 |
| middle temporal gyrus, superior temporal gyrus, temporal pole (superior & middle temporal gyri), inferior temporal gyrus | R | 60 | -34 | 0 | 7.10 | 1117 |
| cuneus, superior occipital gyrus, calcarine fissure | L/R | 14 | -92 | 26 | 5.98 | 334 |
| superior frontal gyrus (dorsolateral & medial) | L | -8 | 54 | 34 | 5.95 | 678 |
| superior frontal gyrus (medial orbital), anterior cingulate cortex (pregenual), superior frontal gyrus (medial) | L/R | -8 | 50 | -8 | 5.33 | 194 |
| insula, putamen, rolandic operculum, inferior frontal gyrus (pars opercularis) | L | -36 | 4 | 6 | 5.09 | 127 |
| angular gyrus, middle temporal gyrus, middle occipital gyrus | L | -40 | -58 | 26 | 4.56 | 167 |
| **DYS (*n* = 21) visual runs (**FWEc = 665) | | | | | | |
| inferior frontal gyrus (pars triangularis, orbitalis, opercularis), posterior orbital gyrus | L | -48 | 24 | 0 | 6.54 | 665 |
| middle temporal gyrus, superior temporal gyrus, supramarginal gyrus | L | -62 | -32 | 2 | 6.40 | 677 |
| **CON (*n* = 29) auditory run** (FWEc = 124) | | | | | | |
| middle temporal gyrus, postcentral gyrus, cingulate gyrus (mid part), superior parietal gyrus, precentral gyrus, precuneus, middle occipital gyrus, supramarginal gyrus, inferior parietal gyrus, inferior temporal gyrus, calcarine fissure, fusiform gyrus, superior temporal gyrus, angular gyrus, temporal pole (superior & temporal gyri), cuneus, supplementary motor area, lingual gyrus, inferior frontal gyrus (pars triangularis, orbitalis), posterior cingulate gyrus, superior frontal gyrus (dorsolateral), inferior occipital gyrus, posterior orbital gyrus, paracentral lobule, rolandic operculum, parahippocampal gyrus, lateral orbital gyrus, middle frontal gyrus, superior occipital gyrus, anterior orbital gyrus | L/R | -52 | -12 | -10 | 9.75 | 13444 |
| middle temporal gyrus, superior temporal gyrus, temporal pole (superior temporal gyrus), inferior temporal gyrus | R | 50 | -4 | -20 | 7.98 | 1038 |
| putamen, insula, rolandic operculum, pallidum, Heschl’s gyrus, amygdala, hippocampus, thalamus (lateral geniculate) | L | -34 | -20 | 8 | 6.15 | 463 |
| thalamus (mediodorsal medial magnocellular, intralaminar, pulvinar medial, ventral posterolateral, mediodorsal lateral parvocellular, ventral lateral, pulvinar anterior, lateral posterior, pulvinar inferior, medial geniculate), hippocampus, lingual gyrus, parahippocampal gyrus | L | -10 | -20 | 0 | 5.82 | 259 |
| superior frontal gyrus (dorsolateral & medial), middle frontal gyrus | L/R | -6, | 60 | 34 | 5.81 | 438 |
| thalamus (ventral posterolateral, pulvinar medial, pulvinar anterior, lateral geniculate, intralaminar, pulvinar inferior, pulvinar lateral, ventral lateral, medial geniculate), hippocampus | R | 14 | -20 | 4 | 5.68 | 149 |
| middle occipital gyrus, middle temporal gyrus, angular gyrus, superior occipital gyrus, superior temporal gyrus, superior parietal gyrus, cuneus, supramarginal gyrus, inferior parietal gyrus | R | 36 | -76 | 40 | 5.46 | 978 |
| cerebellar hemispheres (lobules IV, V, VI, VIII, IX, crus I, crus II), vermis (lobules VI, VII, VIII, IX) | L/R | 12, | -50 | -30 | 5.35 | 630 |
| inferior frontal gyrus (pars triangularis) | L | -50 | 22 | 20 | 4.54 | 124 |
| **DYS (*n* = 21) auditory run** (FWEc = 192) | | | | | | |
| middle temporal gyrus, superior temporal gyrus, temporal pole (superior temporal gyrus) | L | -52 | -12 | -12 | 7.00 | 403 |
| middle temporal gyrus | L | -56 | -38 | 0 | 5.33 | 192 |

*Note*. L – left, R – right. All results are reported at *p* < .001 height threshold corrected for multiple comparisons using *p* < .05 FWE cluster threshold

**Table S6.** *Logical Conjunction Results From* *Paired T-Tests for Both Visual (Words > False Fonts) and Auditory Runs (Words > Words Backward) Within Control (CON) and Dyslexic Groups (DYS)*

| Brain regions | Hemisphere | Peak of cluster coordinates | | | *t-*value | Number of voxels |
| --- | --- | --- | --- | --- | --- | --- |
|  |  | x | y | z |  |  |
| **CON (*n* = 29) visual and auditory runs conjunction** (FWEc = 143) | | | | | | |
| middle temporal gyrus, inferior frontal gyrus (pars triangularis, orbitalis, opercularis), superior temporal gyrus, temporal pole (superior & middle temporal gyri), supramarginal gyrus, posterior orbital gyrus, inferior parietal gyrus, angular gyrus, lateral orbital gyrus, anterior orbital gyrus, middle occipital gyrus | L | -54 | -58 | 12 | 9.58 | 3164 |
| middle temporal gyrus, superior temporal gyrus | R | 50 | -34 | -2 | 7.98 | 186 |
| superior frontal gyrus (dorsolateral & medial) | L | -6 | 56 | 36 | 5.94 | 430 |
| superior temporal gyrus, temporal pole (superior & middle temporal gyri), middle temporal gyrus | R | 52 | 10 | -16 | 5.76 | 175 |
| precuneus, posterior cingulate gyrus, middle cingulate & paracingulate gyri | L/R | -2 | -56 | 30 | 5.48 | 448 |
| supramarginal gyrus, postcentral gyrus, rolandic operculum, superior temporal gyrus | L | -50 | -24 | 22 | 5.35 | 217 |
| middle cingulate & paracingulate gyri | L/R | -2 | -10 | 40 | 5.17 | 143 |
| **DYS (*n* = 21) visual and auditory runs conjunction** (FWEc = 545) | | | | | | |
| middle temporal gyrus, superior temporal gyrus | L | -52 | -12 | -6 | 6.73 | 545 |

*Note*. L – left, R – right. All results are reported at *p* < .001 height threshold corrected for multiple comparisons using *p* < .05 FWE cluster threshold

**Table S7.** *Results* *From the Flexible Factorial Model for Both Visual (Words > False Fonts) and Auditory Runs (Words > Words Backward) Between Control (CON) and Dyslexic Groups (DYS)*

| Brain regions | Hemisphere | Peak of cluster coordinates | | | *t-*value | Number of voxels |
| --- | --- | --- | --- | --- | --- | --- |
|  |  | x | y | z |  |  |
| **CON > DYS main effect of group for visual and auditory runs** (FWEc = 121) | | | | | | |
| middle temporal gyrus, supramarginal gyrus, superior temporal gyrus, inferior temporal gyrus, inferior parietal gyrus, middle occipital gyrus, fusiform gyrus, angular gyrus, postcentral gyrus, rolandic operculum | L | -58 | -30 | 30 | 7.19 | 2181 |
| rolandic operculum, superior temporal gyrus, middle temporal gyrus, supramarginal gyrus, insula, postcentral gyrus, precentral gyrus, inferior frontal gyrus (pars opercularis), putamen, inferior temporal gyrus | R | 46 | -4 | 14 | 7.07 | 1512 |
| superior frontal gyrus (dorsolateral & medial), middle frontal gyrus | R | 22 | 34 | 54 | 6.56 | 464 |
| insula, rolandic operculum, superior temporal gyrus, Heschl’s gyrus | L | -40 | -6 | 18 | 6.21 | 266 |
| middle cingulate & paracingulate gyri, paracentral lobule, supplementary motor area, precuneus | L/R | -8 | -26 | 42 | 5.96 | 1108 |
| precentral gyrus, superior frontal gyrus (dorsolateral) | L | -32 | -8 | 68 | 5.70 | 191 |
| angular gyrus, supramarginal gyrus, inferior parietal gyrus, middle occipital gyrus | R | 52 | -62 | 40 | 5.37 | 516 |
| superior frontal gyrus (dorsolateral and medial), middle frontal gyrus | L | -18 | 38 | 34 | 5.13 | 241 |
| precentral gyrus, postcentral gyrus, middle frontal gyrus | R | 58 | -12 | 46 | 5.09 | 140 |
| lingual gyrus, cerebellar hemisphere (lobule VI), fusiform gyrus | R | 14 | -68 | -10 | 5.04 | 209 |
| temporal pole (superior temporal gyrus), middle temporal gyrus, superior temporal gyrus | L | -54 | 10 | -16 | 5.00 | 156 |
| supplementary motor area, middle cingulate & paracingulate gyri | L/R | 4 | 6 | 48 | 4.93 | 174 |
| postcentral gyrus, superior parietal gyrus | R | 24 | -48 | 58 | 4.85 | 121 |
| postcentral gyrus, supramarginal gyrus | R | 38 | -32 | 36 | 4.78 | 137 |
| **DYS > CON main effect of group for visual and auditory runs** | | | | | | |
| No suprathreshold clusters |  |  |  |  |  |  |

*Note*. L – left, R – right. All results are reported at *p* < .001 height threshold corrected for multiple comparisons using *p* < .05 FWE cluster threshold

**Figure S1.**

*Main Effect of Group CON > DYS for Both Visual (Words > False Fonts) and Auditory Runs (Words > Words Backward)*

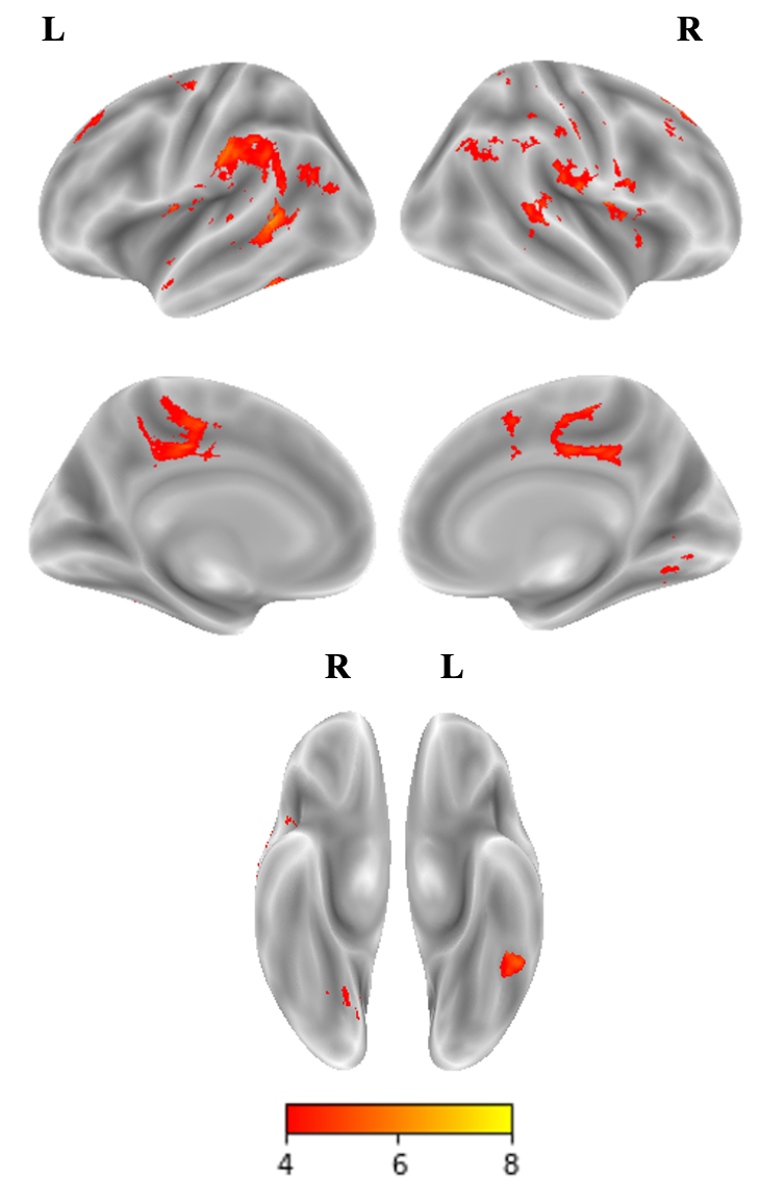

*Note*. CON – control group, DYS – dyslexic group, L – left hemisphere, R – right hemisphere. Results reported at *p* < .001 height threshold corrected for multiple comparisons using *p* < .05 FWE cluster threshold.

**Table S8.** *MRS Checklist*

| **1. Hardware** |  |
| --- | --- |
| a. Field strength [T] | 7T |
| b. Manufacturer | GE Healthcare |
| c. Model (software version if available) | Discovery MR 950 |
| d. RF coils: nuclei (transmit/ receive), number of channels, type, body part | H^1^  32 channel  head coil |
| e. Additional hardware | MR Safe Response PAD |
| **2. Acquisition** |  |
| a. Pulse sequence | Semi-Laser |
| b. Volume of Interest (VOI) locations | left superior temporal sulcus (STS) |
| c. Nominal VOI size [cm^3^, mm^3^] | 15 x 15 x 15 mm^3^ |
| d. Repetition Time (TR), Echo Time (TE) [ms, s] | TR = 4000ms TE = 28ms |
| e. Total number of Excitations or acquisitions per spectrum | 320 averages |
| f. Additional sequence parameters  (spectral width in Hz, number of spectral points, frequency offsets) | 5000 Hz, 4096 points |
| g. Water Suppression Method | VAPOR |
| h. Shimming Method, reference peak, and thresholds for “acceptance of shim” chosen | Automated linear shims adjustment, 0 and first shim order only, water peak, < 20 Hz |
| i. Triggering or motion correction method  (respiratory, peripheral, cardiac triggering, incl. device used and delays) | N/A |
| **3. Data analysis methods and outputs** |  |
| a. Analysis software | fsl-mrs (version 2.0.7) |
| b. Processing steps deviating from quoted reference or product | fsl_mrs default pipeline + simulated basis set |
| c. Output measure  (e.g., absolute concentration, institutional units, ratio) Processing steps deviating from quoted reference or product | Ratio to total creatine |
| d. Quantification references and assumptions, fitting model assumptions | The customized basis set includes: Ala, Asc, Asp, Cit, Cr, EtOH, GABA, GPC, GSH, Glc, Gln, Glu, Gly, Ins, Lac, NAA, NAAG, PCh, PCr, PE, Phenyl, Scyllo, Ser, Tau, Tyros, bHB, bHG. Macromolecules MM09, MM12, MM14, MM17, MM21 were added. |
| **4. Data Quality** |  |
| a. Reported variables  (SNR, Linewidth  (with reference peaks)) | *Linewidth (for metabolite group reported by fsl_mrs): mean 11.11 Hz,  SD = 2.79 Hz SNR (for NAA reported by fsl_mrs): mean 7.05,  SD = 23.95* |
| b. Data exclusion criteria | *Linewidth > 20 Hz, CRLB > 20% and Visual inspection (baseline, residuals)*  *Glu - 4 subjects excluded GABA – 7 subjects excluded* |
| c. Quality measures of postprocessing Model fitting (e.g., CRLB, goodness of fit, SD of residual) | *%CRLB of Glu: mean 2.96, SD = 0.79*  *%CRLB of GABA: mean 10.59, SD = 2.76*  *%CRLB of NAA: 1.76 SD = 0.46* |
| d. Sample Spectrum | 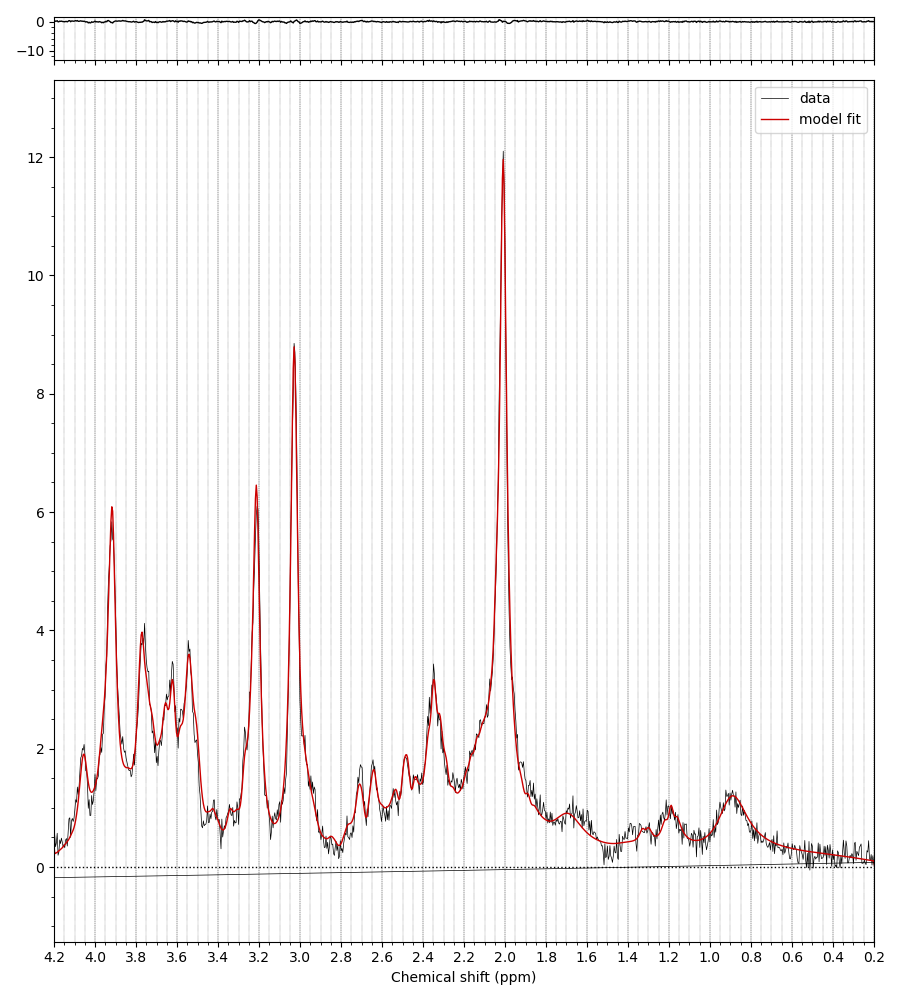 |
